# An optopharmacological tool for on-demand perturbation of branched actin polymerization

**DOI:** 10.64898/2026.05.28.728449

**Authors:** Srishti Mandal, Alka Chahal, Steve Chung, Arka Som, Kamal Kishore, Parinaz A. Shekari, Amit Baidya, Kavita Babu, Michael Mak, Jonathon A. Ditlev, Sarit S. Agasti, Sudha Kumari

## Abstract

Regulated actin cytoskeletal organization and dynamics are essential for life-sustaining processes in eukaryotic cells. The spatiotemporal regulation of filamentous actin can generate a staggering variety of higher-order architectures to meet diverse cellular needs. However, connecting a specific actin substructure to the selective cellular function remains challenging because tools to specifically inhibit F-actin in a spatiotemporally controlled manner are currently limited. To address this crucial technical gap, we have synthesized and characterized a photocrosslinkable small molecule inhibitor of a major and evolutionarily conserved actin nucleation factor, Arp2/3 complex. The new inhibitor “AZ” combines the specificity of a preexisting inhibitor CK666 with photosensitive crosslinking by an Azido group, such that a light-induced, instantaneous, on-demand, irreversible inhibition of Arp2/3 complex-dependent actin nucleation is achieved in a spatiotemporally defined manner at a subcellular, supracellular, and organismal scale. Whole cell treatment with AZ, followed by subcellular photoirradiation, revealed a divergent actin nucleation homeostasis in the cortex of different cells. Thus, AZ could be a powerful tool to infer actin polymerization homeostasis and network properties in live cells-processes which otherwise remain obscure.

## Introduction

Filamentous Actin (F-actin) forms fundamental cytoskeletal scaffolds and architectures in eukaryotic cells, mediating essential cellular processes. Regulated assembly of branched F-actin in space and time is proposed to mediate evolution and patterning of subcellular and supracellular behaviours such as generation and propagation of mechanical forces, compartmentalization of signalling molecules, cell growth and division, and intercellular communication, to name a few. However, spatial and kinetic interrogation of these processes is a challenge. This is because a common pool of actin monomers is utilized and repurposed to generate divergent organizations ^1–3^, and routine pharmacological agents that target actin monomer dynamics lack spatial control on actin assembly, such that all actin organizations are simultaneously affected in a treated cell. In addition, while treatment of cells with pharmacological agents involves a clean temporal start point, their buildup inside the cell can still be slow, sometimes of the order of minutes, limiting a clean temporal handle on the inhibition as well ^4^. Given the spatiotemporal complexity of actin inhibition in such assays, the results are challenging to interpret and do not allow discrimination of whether or how a specific actin organization enables a given cellular behaviour. Furthermore, the cellular actin cytoskeleton is dynamic, and the cytoskeletal states of cells, such as immune cells, evolve rapidly in time, where each cytoskeletal state likely constructs critical biochemical and biophysical make-up, essential for myriad cellular functions ^5–12^. Here, inhibiting actin polymerization in a preceding state can have profound effects on all the subsequent cellular states ^8,13^, limiting mechanistic insights into the role of actin dynamics in a given state. Thus, a controlled ablation of actin polymerization in both time and space is required to establish the structure-function relationship between actin networks and cellular behaviours. While genetic overexpression systems, such as overexpression of light-sensitive protein domains coupled to actin regulatory proteins ^14,15^, have previously been developed for spatial control of cytoskeletal manipulation, the stochasticity and variations in overall cellular levels, as well as a lack of specificity for a particular actin organization (branched vs. linear actin), limit the benefits of these systems. The overexpression systems face further challenges in primary cells since these cells are usually hard to transfect and genetically manipulate. The latter limitation is overcome in light-induced inhibitory photoswitches targeting actin monomers ^16–18^; unfortunately, the readout from photoswitch data is still complex due to reversible chemistry of inhibition, requiring multiple rounds of illumination to achieve effective inhibition, diffusion of both the inhibitor and target molecules in the cytoplasm ^19^, and a lack of specificity for branched vs. linear actin organizations. Thus, a cell-permeable and irreversible agent to precisely inhibit actin polymerization in space and time, without targeting actin monomers directly, is required to gain deeper insights into how a specific actin architecture enables a selective cellular behaviour ^20,21^.

The diversity of F-actin architectures and dynamics within a cell is achieved via selective localization and activation of actin effector proteins, including nucleation-promoting factors (NPFs) and nucleation factors (NFs). The major branched actin nucleating machinery, where most regulatory processes converge, is composed of the Arp2/3 complex—an evolutionarily conserved, seven-subunit complex that is recruited to preexisting filaments to initiate fresh branches. Hypothesizing that controlling the activity of the Arp2/3 complex in space and time will impart control over branched actin dynamics on demand, we sought to generate a cell-permeable, irreversible, optopharmacological inhibitor of the Arp2/3 complex.

The small molecule inhibitor CK666 is the current state-of-the art pharmacological tool for inhibiting branched actin polymerization via Arp2/3 complex ^22–24^. CK666 binds within a pocket in the Arp2/3 complex and stabilizes its splayed conformation, with Arp2 and Arp3 subunits prevented from moving into a short-pitch conformation required for actin dimer loading and filament elongation at a branchpoint ^22–24^. Its use has provided a wealth of information on branched actin involvement in diverse processes including cell migration, adhesion, cell-cell communication, and animal developmental dynamics. CK666 also provides an acute temporal handle on branched actin polymerization, and because it is cell permeable it is also widely used in primary cells. However, its binding is reversible^22,25,26^, and as a diffusible entity, it cannot be used to spatially control the inhibition of the Arp2/3 complex. We reasoned that installing a photocrosslinkable chemical group on CK666 would provide a powerful handle on its activity and allow irreversible and spatiotemporally controlled inhibition of actin polymerization by the Arp2/3 using light. We thus generated Azido-CK666 (“AZ”) by using a novel synthetic scheme. AZ showed a light-induced switch in its inhibitory activity in *in-vitro* actin polymerization assays, and impeded actin-dependent behaviours at subcellular, cellular, and organismal level, on-demand. Furthermore, AZ uncovered the differences in homeostatic actin nucleation state of cell cortex, which was not possible before. Thus, we propose AZ as a tool to interrogate branched actin dependent processes in space and time at diverse scales across many cell biological model systems.

### Conceptualization and design of AZ

The currently available state-of-the-art tool for Arp2/3 complex inhibition is the small molecule inhibitor CK666 that binds non-covalently and reversibly within a conserved hydrophobic pocket between the Arp2 and Arp3 subunits ^22–24^. We leveraged CK666’s structure and the structural insight of the Arp2/3 complex binding pocket to design a light-activated covalent inhibitor—Azido-CK666 (“AZ”). The key to this design was the installation of the Azido group into the structure, which is chemically inert under physiological conditions but is activated by light to form new covalent linkages ^27^. Light irradiation (typically using light in the range from 254 nm up to 405 nm) activates the aryl azides to form a highly reactive nitrene intermediate, which then undergoes X–H insertion (X = C, N, O, and S) or another prominent pathway where it rearranges to generate a highly reactive 7-membered ketenimine heterocycle, which then gets coupled with a biological nucleophile (e.g., an amine from a lysine side chain) ^28,29^ (Figure 1A). Because the nitrene or its rearranged ketenimine intermediate is highly reactive and short-lived, the covalent attachments with the surrounding groups typically occur at close intermolecular distances, making it a highly localized crosslinking agent. Thus, when integrated along with the Arp2/3 complex binding domain of CK666, the Azido group would lead to the covalent anchoring of the inhibitor within the Arp2/3 complex binding pocket, effectively locking the inhibitor in, thereby enhancing pharmacological responses by increasing the inhibitor’s residence time and, importantly, restricting its activity to defined spatial regions, thus enabling precise spatiotemporal control through light irradiation.

**Figure 1:**
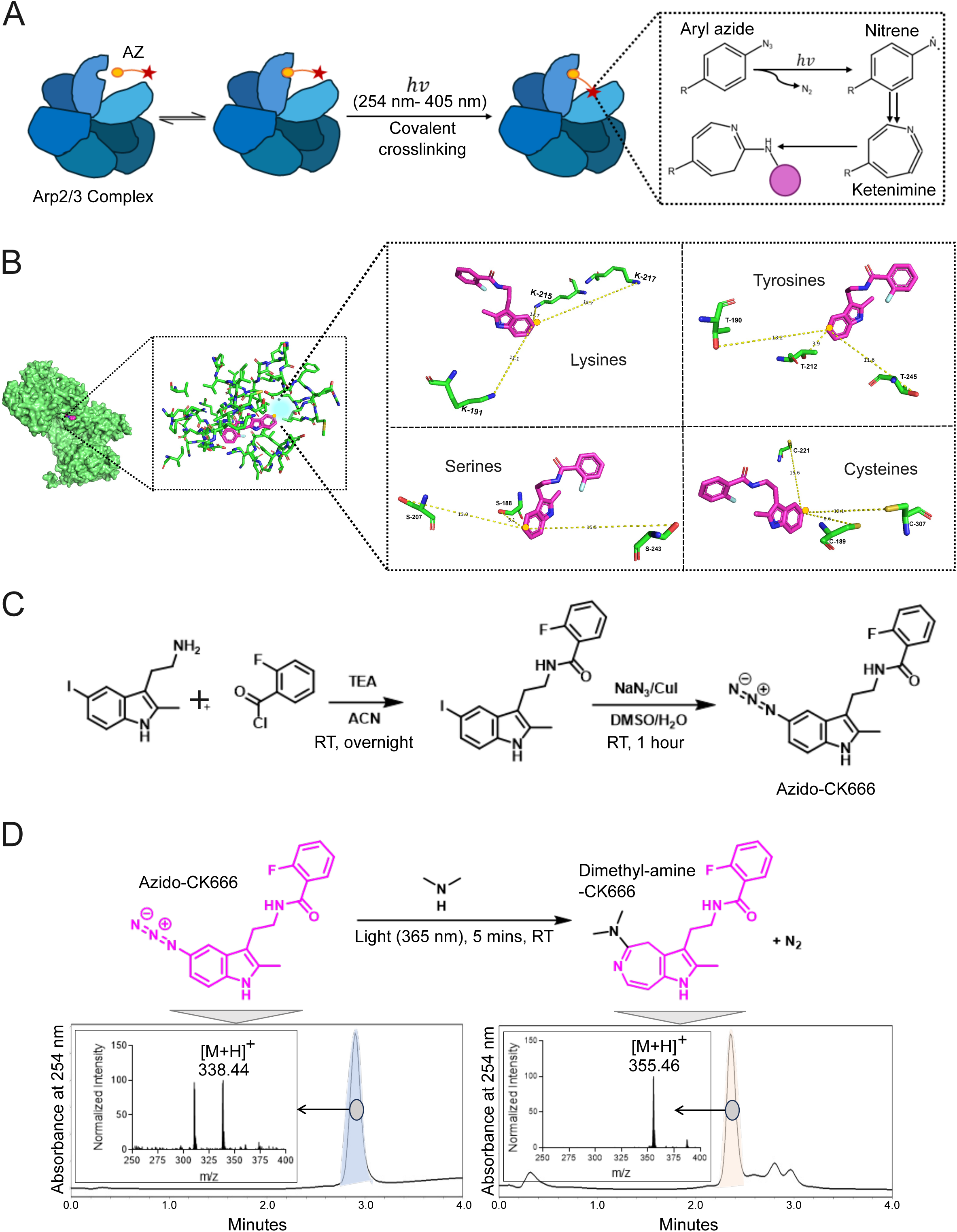
Synthesis and characterization of AZ. (a) A schematic showing the proposed mechanism of Azido-CK666 (AZ) action by covalently crosslinking to the Arp2/3 complex upon photoirradiation. Red star, Azido group; Purple circle, photocrosslinked amino acid in the Arp2/3 complex binding pocket. (b) *In-silico* exploration of photocrosslinkable amino acid residues near CK666 (magenta) with C5 carbon for AZ placement (yellow circle) within the Arp2/3 complex (green) binding pocket (cyan), indicating proximal Lysine, Tyrosine, Serine, and Cysteine residues. (c) Total synthesis scheme of AZ showing the two-step process used to generate the inhibitor. RT- Room Temperature. (d) *In-vitro* photocrosslinking of dimethylamine with AZ in the presence of light (365 nm for 5 minutes), with reactants and products analyzed using LCMS—UV chromatogram (absorbance at 254 nm plotted against time) and intensity plotted against mass/charge peaks (inset graph).

To identify structurally permissive sites for azide installation in CK666 and to gain insight into the accessible crosslinking sites within the binding pocket of the Arp2/3 complex, structural visualization of the vicinity of the Arp2/3 complex protein–inhibitor interaction pocket was performed using PyMOL (Figure 1B). The previously reported crystal structure of the Arp2/3 complex with CK666 (PDB ID: 3UKR) ^23^ was used to guide this rational design and to identify optimal positions for azide installation. We found that the C5 position of the indole ring of CK666 is amenable to modification without causing steric clashes with protein residues or disrupting any crucial protein–inhibitor interaction. Moreover, we found that several potentially nucleophilic residues (including amines from lysine side chains) are within proximity (∼15 Å) to the C5 position of the indole ring of CK666, suggesting a favorable geometry for efficient covalent capture (Figure 1B). Based on these insights, we chose to install the azide group at the C5 position of the indole ring of CK666, to enable proximity-driven crosslinking while retaining its binding potential. To generate AZ, we devised a novel two-step synthesis scheme that used 2-fluorobenzoyl chloride and 2-(5-iodo-2-methyl-1H-indol-3-yl)-ethylamine as starting materials, thereby bypassing the need to install modifications directly onto CK666 and making the synthesis more scalable and cost-effective (Figure 1C). The yield of the reaction was high in both steps. The intermediate and the final compound were characterized, and the purity of compounds at each step of synthesis was estimated using nuclear magnetic resonance (NMR) and high-resolution mass spectrometry (HRMS) to support the successful synthesis of the AZ compound (Supplementary Fig. 1A–D). After successful total synthesis, we validated the photocrosslinking activity of AZ using an *in-vitro* analysis, where dimethylamine was used as a model nucleophile. A mixture of AZ and dimethylamine was irradiated with 365 nm light, and the reaction product was analyzed using LC–MS. We found that photoirradiation yielded a hybrid compound (Dimethylamine–CK666; Figure 1D), covalently linking dimethylamine to the CK666 core. Importantly, the cross-linked product was not observed in the absence of light irradiation (Supplementary Fig. 1E), confirming that *in-vitro* crosslinking is strictly light-dependent. ^18–20^ Together, these results indicated that AZ has the potential to function as a light-activated covalent inhibitor.

### *In-vitro* and *in-cellulo* application of AZ

Our hypothesized mode of AZ-directed inhibition is that when AZ is irradiated with UV light ranging between 254 nm and 405 nm in the presence of the Arp2/3 complex, AZ will bind the Arp2/3 complex between Arp2 and Arp3 subunits and covalently crosslink to surrounding amine-carrying residues. To test if AZ could indeed inhibit the actin nucleating activity of the Arp2/3 complex in a light-dependent manner, we performed an *in vitro* actin polymerization assay ^30^ with purified Arp2/3 complex, unlabeled monomeric actin (G-actin), pyrene-actin, and AZ or CK666 at varying concentrations, in the presence and absence of 365 nm light (Figure 2A, see ‘Methods’). Before initiating actin polymerization assays to test AZ inhibition, mixtures of AZ and Arp2/3 complex were irradiated with UV light for 10 minutes (“+ Light”; Figure 2B), and in the polymerization assays, pyrene fluorescence was measured over time as a readout of actin nucleation-dependent polymerization. A distinct drop in pyrene fluorescence signal was observed at 25 µM and above concentrations of AZ only in photo-irradiated samples, while the non-irradiated samples remained unaffected (Figure 2B). CK666 activity, on the other hand, was independent of light, as the inhibition of polymerization was seen in both irradiated and non-irradiated samples above a threshold concentration, as expected. These data confirmed that above a critical concentration and photoirradiation, AZ inhibits actin nucleation by the Arp2/3 complex.

**Figure 2:**
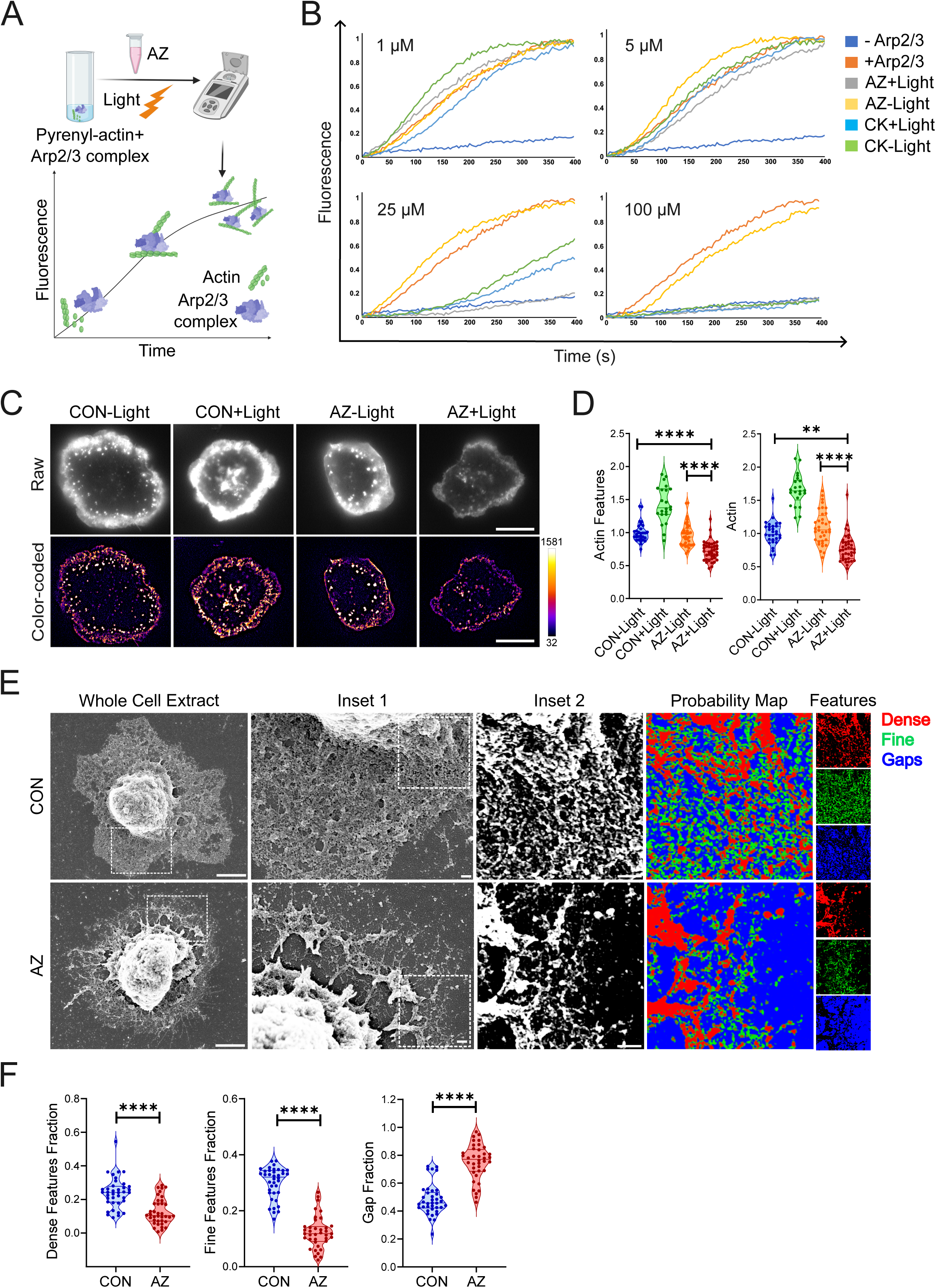
***In-vitro* and *in-cellulo* characterization of AZ activity** (a) Schematic illustrating *in-vitro* actin polymerization assays performed using purified pyrene-actin, Arp2/3 complex, N-WASP, and inhibitors—AZ or CK666 (CK). (b) Plots showing Pyrene fluorescence over time, with varying concentrations of AZ and CK. “+ or - Light” indicates the presence or absence of photoirradiation. N = 2 replicates with similar results. (c) TIRF microscopy images comparing the F-actin levels (phalloidin) at Jurkat T-cell immunological synapses, in raw and color-coded (processed to extract and highlight high-intensity F-actin features using pseudocolor, see ‘Methods’ section). “CON”, untreated control cells; “AZ”, AZ-treated cells;“+ or - Light”, presence or absence of photoirradiation. Scale bar = 10 µm. (d) Plots comparing the mean intensity of F-actin in raw (“Actin”) and processed images (“Actin Features”) across treatment conditions. Each point in the graph represents mean values derived from a single cell, further normalized to “CON-Light”, ± Standard Error of Mean (n= 20-40 cells analyzed per condition). **P ≤ 0.01, ****P ≤ 0.0001, as determined using a Kruskal-Wallis test with Dunn’s multiple comparisons test. N = 3 replicates with similar results. (e) Scanning Electron Microscopy (SEM) images of detergent-extracted Jurkat T-cell immunological synapses. Inset 1 shows a magnified view of the region specified in the image on the left, and Inset 2 is a further magnified zone demarcated in Inset 1 to show finer details of the cytoskeleton. Probability map images show dense features (red), fine features (green), and gaps (blue), corresponding to Inset 2. “CON”, untreated control cells; “AZ”, AZ-treated cells. Left images’ scale bar = 2 µm; Inset 1 and 2 scale bar = 200 nm. (f) Plots comparing the fraction of total pixels occupied by dense features, fine features, or gaps. Each point in the graphs represents mean values derived from a single analyzed area (as described in ‘Methods’) ± Standard Error of Mean (n= 38 areas analyzed per condition). ****P ≤ 0.0001, as determined by a Mann-Whitney U Test. N = 2 replicates with similar results.

To test whether AZ inhibits branched actin polymerization in cellular systems as well, we tested AZ activity in T-cells. In response to activation stimuli presented on antigen presenting cells, T-cells generate specialized cell-cell contact interfaces termed immunological synapses ^31,32^, where highly dynamic Arp2/3 complex-dependent actin architectures are assembled- a process crucial for optimal T-cell activation ^5^. We first determined the safe time limit of a cell’s exposure to blue light in our illumination setup by assaying long-term effects of light exposure on primary T-cell proliferation—a sensitive readout of T cell health ^33^. Up to 2 minutes of exposure to 254 nm light did not yield a measurable block in stimulation-dependent proliferation of murine primary T-cells (Supplementary Fig. 2). We thus chose an intermediate exposure time of 1.5 minutes as a safe light exposure time limit. Then, to observe the acute effect of AZ treatment on T-cells’ actin cytoskeleton, T-cells were suspended in media containing AZ, exposed to 254 nm light for 1.5 minutes, and then allowed to adhere to glass substrates coated with stimulatory and adhesion ligands anti-CD3 and ICAM-1, to form immunological synapses ^34,35^. The level of F-actin (marked by phalloidin) at synapses was quantified using total internal reflection fluorescence (TIRF) microscopy. Cells treated with AZ and irradiated with light (“AZ + Light”) displayed significantly diminished phalloidin levels at synapse, indicating photo-inhibition of F-actin nucleation and polymerization (Figure 2C-D, Supplementary Fig. 3A-B). Consistently, immunofluorescence of Tyr378/397 phosphorylated form of Hematopoietic lineage cell-specific protein 1 (phospho-HS1), an actin branch-point marker in immune cells ^36^, also showed a significant reduction upon AZ + Light treatment (Supplementary Fig. 3A-B). Interestingly, Arp2/3 complex levels at the synaptic interface, marked by Arp2 immunostaining, were not reduced in AZ + Light treatment, indicating that AZ does not inhibit Arp2/3 complex recruitment to the synaptic plane (Supplementary Fig. 3C-D), an effect also observed in global perturbation of activity using CK666 (“CK+Light”; Supplementary Fig. 3C-D).

Since light exposure is expected to induce a covalent linkage between AZ and the Arp2/3 complex, we hypothesized that light exposure would render AZ activity irreversible, i.e., the inhibitory effect would remain even after AZ is washed out. To test this, we washed out AZ after whole-cell AZ treatment and photoirradiation and observed changes in F-actin levels. Indeed, the levels of F-actin did not recover in the AZ-treated cells (Supplementary Fig. 4A-B). As a control, in cells treated with CK666, which is known to be reversible ^22,25,26^, F-actin levels were restored soon after the washout (Supplementary Fig. 4C-D). These results indicate that within the time frame of immunological synapse experiments, brief, single-shot photoirradiation can generate irreversible AZ activity. This broadens the scope of AZ’s usage to study communication between diverse cell-cell collectives and their interactions, particularly where selective inhibition of chosen cells is desired without affecting other cells.

Actin polymerization plays a crucial role in the dynamics and maturation of the immunological synapse ^5,13,35,37^. Particularly, the F-actin-rich dynamic cortical area defined as the “actin ring” ^8^ is required for synapse symmetry and integrin-based adhesion. The F-actin ring is composed of two major radial zones: the lamellipodial region (the outer ∼20% along the synapse radius) ^38^ and lamella (the remaining ∼80% along the synapse radius) (Supplementary Fig. 5A). To assay the differential requirement of Arp2/3 complex between the branched actin-rich annular zones of actin ring, we treated T-cells with 25 µM AZ and light and imaged the network using Structured Illumination super-resolution microscopy. Detailed examination of images showed a depletion of lamellipodia in treated cells (Supplementary Fig. 5B-C) despite no appreciable difference in cell area (Supplementary Fig. 5D). To validate these observations, we detergent extracted the cellular cytoskeleton ^39^ followed by ultrastructural visualization using scanning electron microscopy (SEM). Analysis of SEM images and network characterization showed a near lack of thick peripheral lamellipodial region in AZ-treated cells (Figure 2E-F, Supplementary Fig. 5E-F). We used the supervised machine learning tool Ilastik ^40^ to generate probability maps of the fraction of “Dense features”, “Fine features”, and “Gaps”—namely, larger bright-presenting structures, finer and thus dimmer cytoskeletal branches, and empty regions within the network, respectively. The probability maps showed significantly lower fraction of dense and fine features coupled with higher gaps in AZ-treated cells (Figure 2E-F). In addition, there was an overall reduction in network complexity as seen by the reduction in the density of “Trees” (defined as 4 or more linked branches), branches, and junctions in the remaining network in AZ-treated cells (Supplementary Fig. 5E-F). Thus, AZ treatment revealed a spatially organized Arp2/3 complex activity in the peripheral region of T-cell immune synapses. Taken together, the results demonstrate AZ’s light-dependent activity *in-vitro* and in cells.

### Spatially scalable cellular- and organism-level perturbation using AZ

The use of light as an inducer of crosslinking enables scalable spatial control of inhibition, as the photoirradiation region of interest (ROI) can be specified locally in subcellular regions of a cell, to subsets of cells in a population, or to specific parts of an organism, by simply expanding the ROI. The T-cell immunological synapse is a particularly interesting model system for sub-cellular cytoskeleton studies, as multiple branched actin networks are hypothesized to work in concert as the synapse evolves and matures during T-cell activation ^5,13,6^. Within the first few minutes of synapse formation, the cell establishes a balance of in-plane cytoskeletal forces originating in the actin ring, sustaining a symmetric and stable synapse ^41^. To investigate if branched networks in the actin ring are also required for maintaining synapse symmetry, we performed AZ-based subsynaptic photoinhibition in the actin ring on-stage. Soon after irradiating an ROI within the actin ring, we observed an immediate appearance of long, thin, filopodia-like structures, characteristic of Arp2/3 complex inhibition (Figure 3A, upper panels, Movie 1). This effect was followed by whole-cell symmetry breaking within a minute (Supplementary Fig. 6A, Movie 3). Thus, transient, spatially constrained, branched network destabilization using AZ could transform the entire synaptic morphology, indicating a role of branched networks in scaling of forces for synapse stabilization. ^42–44^ Such localized effects on network morphology and overall cell symmetry were absent in cells irradiated in the actin ring in the absence of AZ (Figure 3A, lower panels, Movie 2), indicating that the effect is specific to the inhibitor and not due to phototoxicity from irradiation.

**Figure 3:**
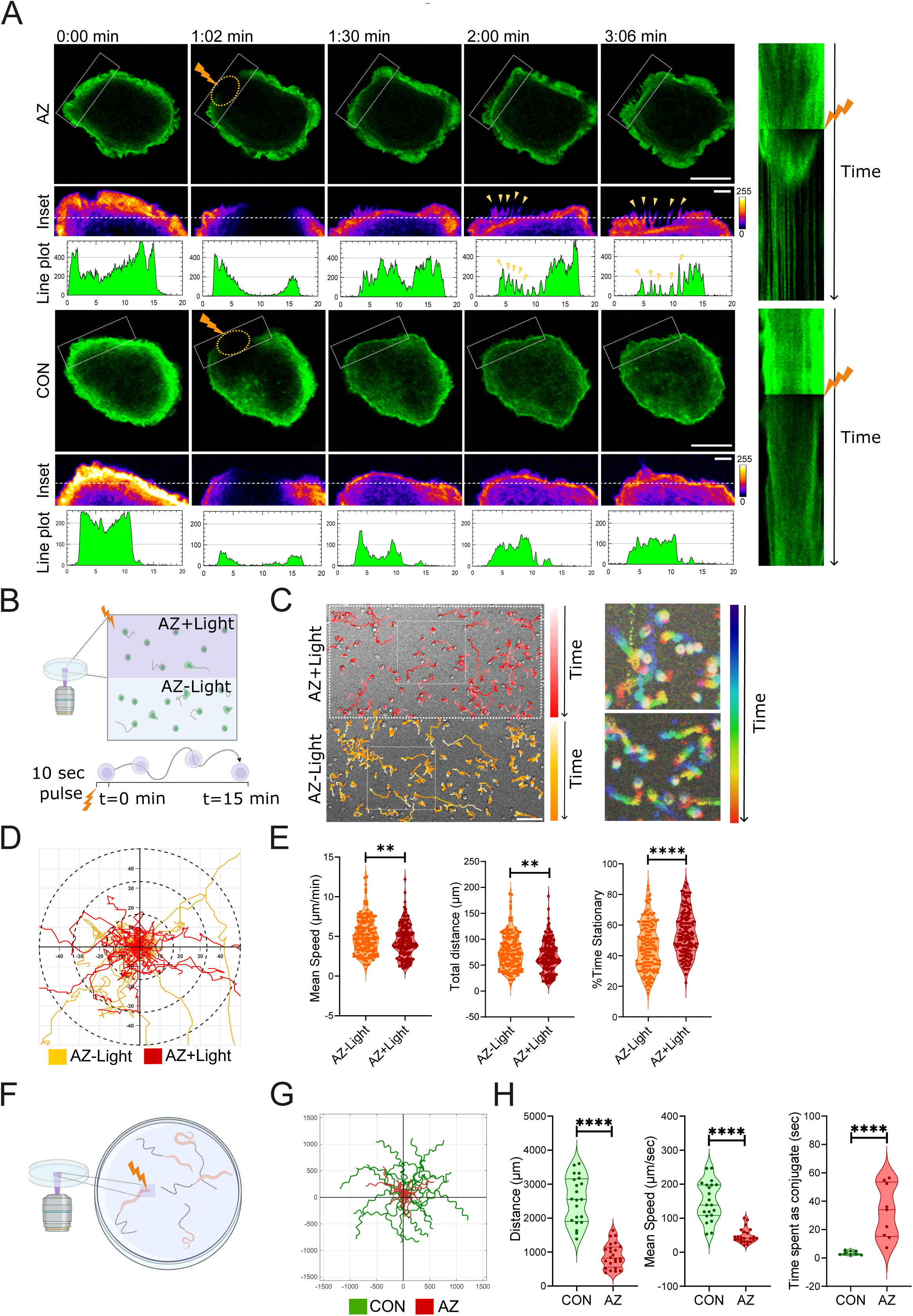
Scalable spatiotemporal perturbation with AZ. (a) Snapshots from a live cell recording of acute localized actin photoinhibition in live Jurkat T-cells expressing Lifeact-citrine. Jurkat T-cell immunological synapses were treated with 5 µM AZ (present in media), followed by on-stage photoirradiation within an ROI in the peripheral actin ring with a 405 nm laser pulse. “AZ”, AZ-treated condition (top panels); “CON”, non-drug control condition (bottom panels). Yellow dotted circle, ROI of irradiation (also marked with lightning bolt); white rectangular box, boundary of magnified insets. Line plots below the insets show actin intensity distribution across the dotted line marked in the insets. Kymographs on the right show changes in local cell shape and curvature upon irradiation in AZ-treated and CON cells. Image scale bar = 10 µm; Inset scale bar = 2 µm. (b) Schematic showing the pattern of acute photoirradiation in a live primary T-cell migration assay, in the presence of AZ. (c) Cell movement trajectories in photoirradiated (“AZ + Light”; top) and non-irradiated (“AZ – Light”; bottom) regions within the same field of view on stage, for a period of 15 minutes. Scale bar = 50 µm. Color bars indicate the progress of the trajectories in each region over time; white squares indicate regions further magnified on the right, showing migration trajectories in a temporal color-code. (d) Trajectories of migrating primary T-cells from photoirradiated and non-irradiated regions. (e) Graphs show quantified cell migration parameters—mean speed, total distance traveled, and percentage of total time spent in the stationary phase. Each point in the graph represents mean values derived from a single cell ± Standard Error of Mean (n= 150-170 cells analyzed per condition). **P ≤ 0.01, ****P ≤ 0.0001, as determined using a Mann-Whitney U Test. N = 2 replicates with similar results. (f) Schematic showing acute organism-level photoinhibition in *C. elegans* using AZ. (g) Trajectories of *C. elegans* movement post-photoirradiation in the presence or absence of AZ, recorded over a period of 15 seconds. “CON”, non-drug control organisms; “AZ”- AZ-treated organisms. (h) Graphs showing total distance travelled by individual *C. elegans* over 15 seconds post-photoirradiation, mean speed during movement, and time spent as a conjugate post-stimulation. Each point in the graph represents mean values derived from a single measurement after up to three photoirradiation pulses ± Standard Error of Mean (n= 10 worms per condition, with 21-24 trajectories for mean speed and distance measurements, and 8-9 conjugate events). ****P ≤ 0.0001, as determined using a Mann-Whitney U Test. N = 2 replicates with similar results.

Light-induced branched actin perturbation can be highly advantageous for population-level investigations, including studies of actin cytoskeletal involvement in cell-cell communication, cell-substrate interaction, or cellular dynamics within live tissues/organisms. In such cases, targeting specific cell/s or region/s within a field of view can yield valuable information on mechanisms of intercellular communication. We leveraged on-stage AZ-based photoinhibition to affect migratory properties of a defined sub-population of primary T-cells, while leaving some cells untouched in the same field of view (Figure 3B). Tracking of cell movement (see ‘Methods’) revealed a change in trajectories with even a short, single photoirradiation event, indicating that dynamic Arp2/3 complex activity regulates long-term migratory capabilities of T-cells, where photoinhibited cells subsequently showed reduced distance travelled and mean speed, and a higher fraction of time spent in the stationary phase (Figure 3C-E, Movie 4). To verify that the observation was not simply an irradiation-induced artifact, the same experiment was performed in the absence of AZ, where the cells did not respond to irradiation (Supplementary Fig. 6B-C).

The spatially defined, on-demand photoperturbation of the actin cytoskeleton using AZ could also be scaled to the organism level. We utilized *Caenorhabditis elegans* as an animal model system to test the application of AZ at the organism level, since migratory responses of *C. elegans* are fast and have been well studied previously ^45–47^, even though underlying cytoskeletal mechanisms are not fully elucidated. For this, age-synchronized L4-stage *C. elegans* were placed on agar plates containing AZ, irradiated with 405 nm laser light, and acute motility responses to the laser pulse were recorded (Figure 3F). While worms in control plates reacted instantly to photoirradiation by moving away rapidly, worms in AZ plates moved significantly more slowly and to shorter distances post irradiation (Figure 3G-H, Supplementary Fig. 6D, Movie 5). Interestingly, occasionally paired worms in control plates separated instantaneously upon photoirradiation. In contrast, paired worms on AZ plates remained conjugated for prolonged periods (Third graph; Figure 3H, Movie 6). The overall reduction in motility of worms upon single-shot photoirradiation on AZ plates continued at least up to after a one-hour rest period (“After Rest”; Supplementary Fig. 6D), highlighting long-term consequences of the short-term AZ-based photoirradiation at the organism level. The slowing down of worms on AZ plates was not a light or AZ drug-related toxicity, since acute effects on motility were not observed if either AZ was missing (“Control”; Supplementary Fig. 6D) or light was missing (“AZ-Light”; Supplementary Fig. 6E). Overall, these data indicated that single-shot, short-term photocrosslinking of AZ can trigger acute as well as long-term effects on worm motility patterns, and the drug could potentially be applied to elucidate mechanisms underlying complex, animal level motility behaviors.

### Inferring subcellular branched actin homeostasis in different cells using AZ

While *in-vitro* reconstituted systems of actin polymerization have been instrumental in visualizing and quantifying the nucleation process and the role of actin effectors in it, tracking local F-actin nucleation dynamics within live intact cells still remains challenging ^48^. We hypothesized that localized perturbation of the Arp2/3 complex using AZ will allow us to gain insights into branched actin nucleation and polymerization dynamics within desired regions of cells. To pursue this, we utilized three cell lines—Jurkat T-cells, Chinese Hamster Ovary (CHO cells), and Human Embryonic Kidney 293T cells (HEK293T): three cell lines used extensively in cell biology research. Cells were transduced with F-Tractin-mCherry, a fluorescent reporter of polymerized actin ^49,50^, to achieve a low-grade expression for efficient F-actin labelling contrast, and were then bathed in media containing 5 µM AZ and imaged using time-lapse imaging. While imaging, cells were irradiated using a 405 nm laser pulse at selected ROIs in their cortical lamellar regions (Figure 4A, Movie 7). A no-drug condition was included as a control (Supplementary Fig. 7A, Movie 8). Photoirradiation in this experimental setup causes both AZ crosslinking as well as an immediate bleaching of the mCherry signal within the ROI. We then recorded the recovery of F-Tractin-mCherry signal within the ROI, plotted, and analyzed it to calculate kinetics parameters t_1/2_ and extent of recovery (“plateau”) (Figure 4B, upper panel; Supplementary Fig. 7E). We found that the F-Tractin recovery profile in the ROI were distinct between the three cell lines, and while Jurkat and HEK293T cells showed marginal to no effect of AZ treatment on recovery parameters (left and right-most graphs; Figure 4B), AZ-treated CHO cells showed significantly lower plateau of recovery compared to control (middle graph; Figure 4B). To explore the underlying causes of the difference in recovery profiles, we measured F-Tractin intensity at whole cell level upon global Arp2/3 complex inhibition via CK666 treatment (bottom graph; Supplementary Fig. 7B). In this assay while both Jurkat ^43^ and CHO cells showed depletion of F-Tractin signal, HEK cells showed a lack of effect, indicating that actin nucleation is insensitive to CK666 in these cells contributing to a lack of effect in photoinhibition assay (top graph; Supplementary Fig. 7B). Considering the differences in recovery between Jurkat and CHO, we reasoned that the signal recovery in the ROI would be majorly influenced by three factors—local Arp2/3 complex activity (expected target of AZ), local Arp2/3 complex diffusion rates, and actin monomer diffusion rates in the cell. Assuming that, due to its small size, actin monomer diffusion is not limited in the cell lines, we did not explore this factor as a determinant influencing F-Tractin recovery. We thus compared the Arp2/3 complex diffusion rates between the cell lines using Fluorescence Recovery After Photobleaching (FRAP) analysis of Arp3-eGFP (Supplementary Fig. 7C-D, Movie 9). We found Arp3 recovery rates to be slower than control F-Tractin recovery rates in CHO cells (≥ 2-fold slower) (Supplementary Fig. 7E), indicating that Arp2/3 complex activity and not the diffusion would be the primary driver of the F-Tractin signal recovery differences in CHOs. Thus, CHO cells strongly rely on Arp2/3 complex activity for cortical lamellar homeostasis. In Jurkats, Arp3 recovery rates was comparable to F-Tractin recovery (Supplementary Fig. 7E) indicating a rapid cytoplasmic flux of Arp2/3 complex would dominate the rate of local F-Tractin recovery in these cells.

**Figure 4:**
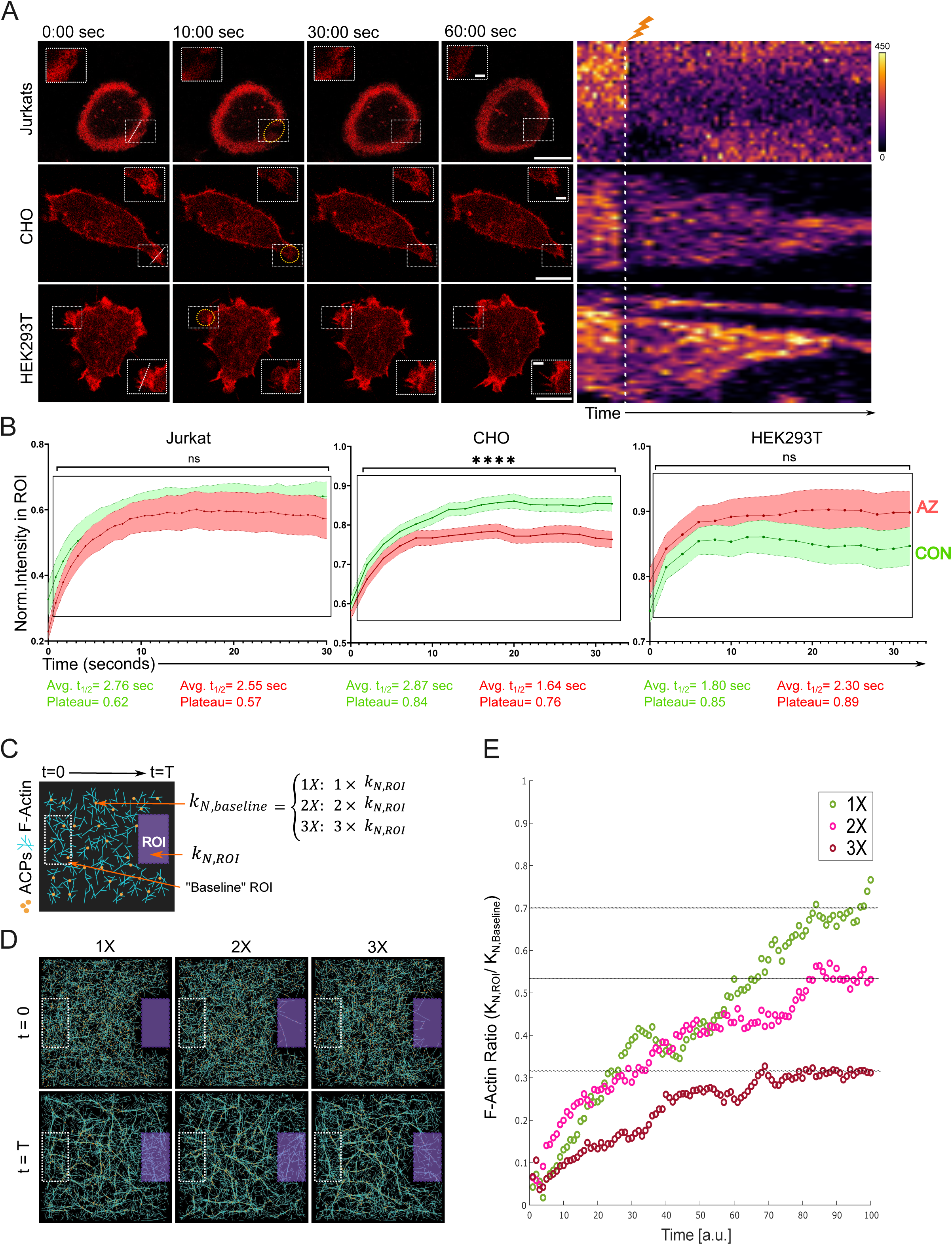
Inferring the subcellular actin nucleation homeostasis in different cell lines. (a) Snapshots from live recordings of acute, localized actin photoinhibition in Jurkat T-cells forming an immunological synapse (upper panel), CHO cells (middle panel), and HEK293T cells (lower panel) expressing F-Tractin-mCherry. Yellow circle, ROI used for photoirradiation and measurement of fluorescence recovery post irradiation; white squares, regions magnified in the insets. Image scale bar = 10 µm; Inset scale bar = 2 µm. Kymographs on the right, generated from the slanted dotted lines in the leftmost snapshots, show the local variation in F-Tractin signal over time. (b) Fluorescence recovery of F-actin (F-Tractin-mCherry) signal in ROI, plotted over time for Jurkats, CHO, and HEK293T cells. Graphs show normalized fluorescence recovery within ROI in untreated cells (“CON”; Green curve), or AZ-treated cells (“AZ”; Red curve). Each curve in the graphs represents the average curve derived from mean intensity values from a single ROI, with normalization to pre-bleach values ± Standard Error of Mean (1 ROI = 1 curve. n= 20-30 cells per condition). ^ns^P ≥ 0.05, ****P ≤ 0.0001, as determined using a Two-way repeated measures ANOVA with Dunnett’s post-hoc multiple comparisons test. N = 3 replicates with similar results. Values below each graph (Green, Control; Red, AZ) indicate the average half-life of recovery (t_1/2_) and the plateau value of the curves. (c) A schematic representation of the computational simulation space showing F-actin filaments (cyan) and actin-crosslinking proteins or ACPs (orange dots), and a region representing localized photoinhibition (“ROI”; purple rectangle) and a representative region of the surrounding non-inhibited space (“Baseline ROI”; dotted rectangle). The ratios describing the specified nucleation rates within the system (ROI and lamellar surround) are denoted on the right. (d) Representative snapshots of F-actin distribution from the start (t = 0) and end (t = T) of the simulations under conditions where baseline nucleation (k_N,Baseline_) varies as 1X, 2X, and 3X k_N,ROI_. (e) Traces of F-actin network recovery in the ROI as quantified from the computational simulation. Dotted lines pass through approximate plateau values of the traces.

To further exploit the F-Tractin recovery profile for understanding lamellar actin homeostasis in CHO cells, we performed computational simulations of actin polymerization in photo-inhibited ROI and compared the resultant trends with experimentally observed traces. In the simulation, we spatially patterned actin nucleation rate (k_N_) within a photoirradiated ROI (k_N, ROI_) and in the surrounding lamellar zone (k_N, Baseline_) (Figure 4C). To simulate AZ treatment, actin filaments within the ROI were depleted at the beginning of simulation, and the nucleation rate was defined as k_N, ROI_; the (k_N, baseline_) of simulation space surrounding the ROI was varied as 1-, 2-, or 3-times of k_N, ROI_, termed “1X”, “2X”, and “3X” respectively (Figure 4C). The 1X case was included to represent the untreated control scenario, since it implies a lack of Arp2/3 complex inhibition. To assess the lamellar nucleation homeostasis by examining the interaction between the ROI and lamellar surround, we plotted the ratio between the actin intensity in the ROI and a region in the lamellar surround against time as the simulation proceeded (Figure 4D-E, Movie 10). The plot showed that the F-actin ratio recovered slower and toward a lower plateau in both 2X and 3X cases, where the profiles of 2X case matched remarkably well with the CHO experimental trace (Control/AZ plateau ratio of experimental ∼1.1 and 1X:2X plateau ratio in simulation ∼1.2), indicating a strong dependence of CHO cell lamellar dynamics on Arp2/3 complex- where the loss of Arp2/3 complex activity in a sub-lamellar zone leads to a ∼50% reduction in polymerization rate compared to surrounding regions- a loss that cannot be compensated by other nucleation factors. Altogether, the rate of signal recovery in an ROI normalized to its local surroundings can be used to map the subcellular Arp2/3 complex activity sustaining subcellular actin network homeostasis in divergent cell types and states.

## Conclusions

Eukaryotic actin cytoskeleton exists in a diversity of forms and dynamics, enabling divergent functions. A major contributor to this diversity is the actin nucleation factor Arp2/3 complex—an evolutionarily conserved multi-subunit assembly ^51^. The Arp2/3 complex is ubiquitously utilized in eukaryotes to generate a staggering diversity of architectures ^21^, which are hypothesized to perform a multitude of functions, most of which remain directly untested due to a lack of tools to inhibit its activity in space and time. Here, we report a novel tool that can perturb Arp2/3 complex activity in a spatiotemporally defined fashion using light, thereby enabling actin structure-function studies. We selected the Arp2/3 complex as the target molecule to interrogate actin dynamics since it is one of the most conserved actin regulators, evolving even before the last common eukaryotic ancestor ^51,52^, and chose light as an inducer for inhibition, as photoirradiation of a sample is easily controllable in time and space using routine confocal microscopes, thus offering an accessible way for precise manipulation.

Spatiotemporal control of actin polymerization has previously been attempted using azobenzene-based photoswitches coupled with actin and microtubule perturbing drugs ^17,18^. Such approach while illuminating for studying the cell biology of cytoskeleton, is limited on two fronts, one, it affects a common pool of monomers thereby simultaneously affecting all organizations, two, exposure to light does not link the inhibitor covalently to the target molecules, and as a result generates highly complex inhibition phenotypes arising from composite effects of the efficacy of ablation, target diffusion, and activity of the target molecule. Use of the Azido group to generate a photocrosslinked inhibitor permanently eliminates the activity of the target molecule, thereby generating a cleaner perturbation readout. This approach could be highly advantageous in highly dynamic cellular and organismal systems where a brief exposure to blue light would acutely inhibit Arp2/3 complex activity, or in multicellular systems where activity in one cell type can be inhibited without affecting its neighbors, since the cross-linked inhibitor will not diffuse out of the target cell. Unlike photoswitches, Azido-based cross-linking is not wavelength-specific and therefore does not require a specialized illumination setup with a UV laser. It only uses low-wavelength light and could be used with 405 nm lasers in routine confocal microscopes ^28,53^. While one brief exposure to 405 nm light was sufficient in most of our assays, the duration of light exposure for photocrosslinking in diverse cellular and organismal systems will require optimization, and for longer-term studies, multiple sequential exposures may be necessary to achieve effective inhibition.

In summary, we provide an optopharmacological tool, generated via total synthesis, for irreversible, multiscale, and spatiotemporal perturbation of actin nucleation events. One limitation of our strategy is that some isoforms of Arp2/3 complex subunits are not CK666 sensitive^54^. This could also be the underlying reason for a lack of inhibition seen in HEK cells (Figure 4). For better coverage of Arp2/3 complex inhibition, our strategy of using the Azido group can also readily be extended to other actin nucleation inhibitors (such as CK869), to achieve more complete Arp2/3 complex inhibition (data not shown) ^24,54,55^. Overall, light-induced Arp2/3 complex inhibition will lead to deeper insights into how cytoskeletal polymers are assembled in a variety of ways to achieve diverse functions at subcellular, cellular, and organismal scales.

## Methods

### Synthesis of Azido-CK666

Part 1: 2-(5-Iodo-2-methyl-1H-indol-3-yl)-ethylamine (0.2 mmol) (Oakwood Chemical) was dissolved in acetonitrile (5 mL) with NEt_3_ (120 µL) and stirred in an ice bath. 2-Fluorobenzoyl Chloride (0.25 mmol) (TCI) was added, and the reaction mixture was stirred in an ice bath for 1 hour, then stirred at room temperature (RT) overnight. H_2_O (30 mL) was added, and the resulting solution was partitioned with ethyl acetate (100 mL). The organic phase was washed with saturated aqueous NaCl and dried with MgSO_4_. The solvent was removed by a rotary evaporator to yield the product (82%). HRMS (ESI-MS) calculated 423.0364 [M+H]^+^, observed 423.0363 [M+H]^+^; calculated 445.01864 [M+Na]^+^, observed 445.0177 [M+Na]^+^.

Part 2: 2-fluoro-N-(2-(5-iodo-2-methyl-1H-indol-3-yl) ethyl) benzamide (0.15 mmol) was dissolved in 2 mL of DMSO: H_2_O mixture (5:1 vol/vol) and pipetted into a 25 mL reaction vessel containing sodium azide (3 mmol), copper(I) iodide (20 mg), and sodium ascorbate (20 mg). N,N’-dimethylethylenediamine (20 µL) was added, and the reaction mixture was stirred for 1 hour at room temperature. The reaction mixture was transferred into 150 mL of ethyl acetate and washed with HCl (0.5 M, 50 mL), H_2_O (50 mL), NaHCO_3_ (sat. 50 mL), H_2_O (50 mL), and NaCl (sat. 50 mL). The organic phase was dried by MgSO_4_ and removed by a rotary evaporator, then dried by vacuum overnight to yield the product, Azido-CK666 (AZ) (92%). HRMS (ESI) *m*/*z*: [M+H]^+^ calculated for C_18_H_17_FN_5_O^+^ 338.1412; observed m/z 338.1414 [M+H]^+^; calculated 360.1232 [M+Na]^+^, observed 360.1229 [M+Na]^+^; calculated 310.1356 [M-N_2_+H]^+^, observed 310.1349 [M-N_2_+H]^+^.

### Structural visualization

PyMOL software was used to visualize all molecular interactions for protein-inhibitor binding. AZ interaction within the Arp2-Arp3 binding pocket was simulated based on the *Bos taurus* Arp2/3 complex and CK666 crystal structure (PDB ID: 3UKR) determined by ^23^. The amino acid residues in closest proximity to the Azido group of the bound inhibitor, which may serve as potential reactive groups in photo-crosslinking reactions, were displayed at up to 15 Å maximum interaction distance.

### *In-vitro* crosslinking

Photochemical conjugation reactions to test the light-induced AZ reactivity towards dimethyl amine were performed in a microcentrifuge tube using a 365 nm ultraviolet light-emitting diode (LED). A reaction mixture was prepared in a microcentrifuge tube by combining AZ (500 µM, from DMSO stock) and dimethylamine (10 mM, from aqueous stock) in Milli-Q water. Subsequently, a photochemical reaction was initiated, wherein the reaction mixture was directly irradiated from the top using the handheld LED light source. The LED intensity was adjusted using a digital UV-LED controller (Thorlabs supplied 365 nm UV LED System (CS2010)), where 100% corresponded to a power/irradiance of ∼ 100 mW/cm^2^ at 365 nm. The irradiation was carried out with approximately 50 mW/cm² emitted light from the handheld LED source for 5 minutes at RT. Following irradiation, a 20 µL solution was injected into the LCMS for analysis of the photochemical reaction outcome.

### Mass spectrometry and NMR

^1^H-NMR spectra were recorded using a Bruker ADVANCE III 400 MHz instrument. Data analysis was done using TopSpin 3.5pl7 and SpinWorks 4. High-resolution mass spectrometry (HRMS) was carried out using Agilent 6538 Ultra High Definition (UHD) Accurate-Mass Q-TOF liquid chromatography-mass spectrometry (LCMS). LCMS for the photochemical crosslinking experiment, monitored at an absorbance of 254 nm, was carried out using a Waters instrument. The solvents used as eluents in LCMS were solvent A (water containing 0.1% formic acid) and solvent B (acetonitrile containing 0.1% formic acid).

### *In-vitro* actin polymerization assay

Non-labelled and pyrene-labelled G-Actin ^30^, Arp2/3 complex ^56^, and N-WASP ^57^ were purified as previously described. 5 µM N-WASP, 1 µM Arp2/3 complex, and 10 µM, 50 µM, 250 µM, 1000 µM AZ or CK666 were mixed in solution and incubated in 365 nm UV light (+ Light samples) or in the dark (- Light samples) for 10 minutes at room temperature. During this incubation, two mixtures were prepared for polymerization reactions: Mix A contained 7.2 µM G-actin, 0.8 µM pyrene-G-actin, 1 mM EGTA pH 8.0, and 100 µM MgCl_2_; Mix P contained 100 mM KCl, 20 mM imidazole pH 7.0, 2 mM MgCl_2_, 2 mM EGTA pH 8.0, and 20 µM ATP. Upon completion of the 10-minute light exposure, N-WASP, Arp2/3 complex, and AZ or CK666 mixture was added to Mix A. The combined mixture was added to Mix P in a quartz cuvette. The final concentrations for each component in the polymerization mixture were 500 nM N-WASP, 100 nM Arp2/3 complex, 3.6 µM non-labelled G-actin, 0.4 µM pyrene-G-actin (4 µM total actin), 50 mM KCl, 10 nM imidazole pH 7.0, 1 mM MgCl_2_, 1 mM EGTA pH 8.0, 10 µM ATP, and 1 µM, 5 µM, 25 µM, or 100 µM AZ, or 1 µM, 5 µM, 25 µM, or 100 µM CK666, as applicable, The quartz cuvette containing the polymerization mixture was placed in a QuantaMaster 80 Spectrofluorometer (Photon Technology International) and actin polymerization was monitored using an excitation wavelength of 365 nm and measuring pyrene fluorescence at its 407 nm emission wavelength for 400 seconds at 4 second intervals at room temperature.

### Mice

C57BL/6J mice used in this study were housed and maintained in the Central Animal Facility (CAF) at the Indian Institute of Science. Dissections were carried out in the lab in a biosafety hood, and all waste was disposed of as per safety regulations.

### Cell culture

Mouse primary T-cells and Jurkat cells were cultured in complete RPMI—RPMI 1640 liquid media (Gibco), with 10% FBS (Gibco), 1 mM Sodium Pyruvate (Gibco), 50 µM β-mercaptoethanol (Gibco), and 1X Penicillin-Streptomycin (Pen-Strep) (Gibco). Adherent cell lines—HEK293T and CHO-AA8—were cultured in complete DMEM—DMEM liquid media (Gibco), with 10% FBS (Gibco) and 1X Pen-Strep (Gibco). Adherent cells were passaged at 70%-80% confluency using Trypsin-EDTA (Gibco). Live imaging was performed in “imaging media”—Phenol Red-free RPMI or DMEM (Gibco) with 10% FBS and 1X Pen-Strep. All media and buffers used were filtered using 0.2 µm filter membranes and stored at 4 °C. All cells were maintained under sterile conditions in a CO_2_ incubator (Eppendorf) at 37 °C and 5% CO_2_.

### Transfections, lentivirus production, and transductions

Plasmid DNA was purified from *E. coli (Stbl3)* bacterial cultures using midiprep kits (Qiagen), and the purity of the DNA was assessed using absorbance ratios (Nanodrop) and agarose gels. Transfections to introduce Arp3-eGFP (pEGFP-N1-ACTR3: Addgene plasmid #8462) into adherent lines were performed using 25 kDa branched PEI (Sigma) with a 3:1 ratio of 1 mg/mL PEI to plasmid DNA, complexed in OptiMEM (Gibco). To introduce the same plasmid into Jurkat T-cells, nucleofection was performed. 1 million Jurkats were resuspended in 100 µL pre-warmed nucleofection buffer (Sodium Phosphate buffer pH 7.2, 1 M KCl, 1 M MgCl_2_, 0.2 M Hepes, 0.24 M Succinic acid, sterile filtered and stored at 4 °C) and mixed with 5 µg plasmid DNA. Nucleofection of this cell suspension was carried out in a 2 mm electroporation cuvette (Biorad) using protocol U-014 (Amaxa Nucleofector II) before gently transferring cells into 1 mL of pre-warmed complete RPMI with 20 U/mL IL-2 (Peprotech) in a 12-well dish and incubating overnight at 37 °C. For lentivirus production of F-Tractin-mCherry (pLV-Ftractin-mCherry: Addgene plasmid #85131), 10 cm dishes of HEK293T cells at 70-80% confluency were transfected using PEI and a total of 10 µg DNA per 10 cm dish, with a DNA ratio as follows: Vector with gene of interest: pMD2.G: psPAX2= 2:1:2. Viral supernatant was harvested at 48- and 72-hours post-transfection. The supernatant was centrifuged, and filtered through 0.45 µm syringe filters before concentration at 4 °C using centrifugal concentrators (Thermo Scientific Pierce Protein Concentrator PES 10K MWCO). Concentrated lentivirus aliquots were stored at -80 °C until use. Lentiviral transductions were performed in adherent cell lines in 6-well dishes—cells were pre-conditioned with 10 µg/mL Polybrene (Sigma) for 30 minutes, followed by the addition of lentivirus into the well and incubation at 37 °C for up to 24 hours, before a media change. Lentiviral transduction in suspension cells (Jurkats) was performed using the spinoculation method. Jurkat cells at 1.5-2 million cells/mL density in a 6-well plate were pre-conditioned with 10 µg/mL Polybrene for 30 minutes, followed by the addition of lentivirus into the well. The plate was then centrifuged at 2000 RCF for 90 minutes at 32 °C, before transferring to 37 °C in the CO_2_ incubator for overnight incubation. Cells were subsequently transferred into fresh RPMI and used in live imaging experiments after 72 hours of virus introduction.

### Primary T-cell isolation

Mice were anesthetized using isoflurane followed by euthanasia by cervical dislocation, and the spleen and lymph nodes (inguinal, mesenteric, axillary, and brachial) were removed. The organs were crushed and strained through a 40 µm strainer into ice-cold RPMI. Cells were centrifuged into a pellet at 600 RCF for 5 minutes, at 4 °C, followed by lysis for 7 minutes on ice using ice-cold 1X RBC lysis buffer (8.02 g NH_4_Cl, 0.84 g NaHCO_3_, 0.37 g EDTA disodium salt, in 100 mL Milli-Q water, for a 10X stock solution, sterile-filtered), then neutralized with PBS. The resulting RBC-free pellet was resuspended in 3 mL warmed complete RPMI and added to a flask pre-coated with 100 µg/mL Goat anti-Mouse IgG (H+L) (Jackson ImmunoResearch Labs Inc). Two rounds of T-cell enrichment (“panning”) were carried out for 40 minutes each, followed by cell counting using a haemocytometer. T-cells were resuspended in complete RPMI with 10 U/mL IL-2 (Peprotech) and stimulated with PMA (25 ng/mL) (Sigma) and Ionomycin (0.5 µM) (Sigma) overnight, followed by transfer into complete RPMI with 10U/mL IL-2, before using in immunocytochemistry and migration assays.

### CFSE proliferation assay and flow cytometry

Panned mouse primary T-cells were washed with PBS and resuspended in 1 µM CellTrace CFSE (Thermo) solution made in PBS. Cells were incubated on a rocker at RT for 20 minutes, protected from light. Complete RPMI was added to quench the reaction, and the tube was incubated at room temperature for 5 minutes. Cells were resuspended in complete RPMI containing 10 U/mL IL-2 and the stimulatory reagents PMA (25 ng/mL) and Ionomycin (0.5 µM). Cells were then photoirradiated in a 35 mm cell culture dish with a transilluminator which irradiates from below (254 nm light, average irradiance = 1.53 mW/cm^2^) for varying times (0.5, 1, 2, 3, 4, and 5 minutes) and incubated at 20,000 cells/well in a 96-well U-bottom plate (NEST) for 48 hours. Cells were stained with live/dead near-IR viability dye (Thermo), fixed in 3.7% PFA, and stored in flow cytometry buffer (1% BSA, 0.5 mM EDTA, 0.1% Sodium Azide, in PBS, sterile-filtered) at 4 °C until acquisition using a BD FACSVerse flow cytometer. Data was analyzed using FlowJo software—the percent of cells divided was inferred using the proliferation modelling function.

### Coverslip-based fixed-cell fluorescence imaging assays

8-well chambers with pre-attached coverslips (Labtek, Thermo) or 18-well chambers (Ibidi) attached with coverslips (#1.5H, 170 µm +/- 5 µm) (Ibidi) were coated at 37 °C overnight with His-ICAM-1 (1 µg/mL) (Sinobiological) and anti-CD3 (5 µg/mL) functional antibody (clone 2C11—mouse or clone OKT3—human) (BioXCell). Coated chambers were washed twice with sterile PBS before use. For TIRF and SIM sample preparation, T-cells (either Jurkats or mouse primary T-cells), in RPMI containing AZ (25 µM, 50 µM, or 100 µM) or the controls (CK666, DMSO) were photoirradiated for 1.5 minutes in a 35 mm dish using a transilluminator (254 nm, irradiance = 1.53 mW/cm^2^). Cells were allowed to form immunological synapses in the coated chambers for 5 minutes at 37 °C, followed by PBS washes, fixation (20 minutes at RT with 3.7% PFA in pre-warmed 1X PHEM buffer (18.14 g PIPES, 6.5 g HEPES, 3.8 g EGTA, 0.99 g MgSO4, pH 7.0 w/ KOH in 1 L water)), permeabilization (3 minutes at RT with 0.1% Triton in PBS), blocking (1 hour at RT with 1% BSA in PBS), and antibody staining. For testing reversibility, cells were washed with PBS or RPMI after irradiation to remove drugs. Primary antibody staining was performed overnight at 4 °C, followed by extensive PBS washes, and secondary staining for 2 hours at RT, also followed by PBS washes. Primary and secondary staining cocktails were made in the blocking solution. Primary staining included Arp2 mAb (clone D85D5, CST) (1:400), pHS1 mAb Tyr378/397 (clone D12C1, CST) (1:800), and the F-actin marker phalloidin-Alexa Fluor 488 (Thermo) (1:500), and secondary staining included Donkey anti-Rabbit IgG (H+L) Rhodamine Red and Donkey anti-Rabbit IgG (H+L) Alexa Fluor 647 (Jackson ImmunoResearch Labs Inc).

### Scanning Electron Microscopy (SEM) of the cytoskeleton

12 mm round coverslips were plasma treated (Harrick plasma cleaner) and coated overnight with His-ICAM-1 (1.5 µg/mL) (Sinobiological) and anti-CD3 functional antibody (5 µg/mL) (clone OKT3, BioXCell). Coverslips were washed twice with PBS before cell addition. Cells treated with 25 µM AZ or DMSO control, and photoirradiation, as previously described for fluorescence imaging assays, were allowed to form immunological synapses on the coverslips for 5 minutes at 37 °C, followed by two PBS washes to remove unbound cells. Samples were processed for cytoskeleton imaging using SEM by a protocol adapted from ^39^. Briefly, cells were incubated with extraction buffer (1% Trition-X-100, 4% 8kDa PEG, 2 µM phalloidin, 2 µM Taxol) for 5 minutes at RT, followed by 3 washes with PEM buffer (100 mM PIPES, 1 mM MgCl_2_, 1 mM EGTA, adjusted to pH 6.9). Extracted cytoskeleton was then fixed with Solution 1 (0.1 M Sodium cacodylate and 2% EM grade glutaraldehyde, in 1X PBS, adjusted to pH 7.3) for 20 minutes at RT, followed by Solution 2 (0.1% Tannic acid in DI water) for 20 minutes at RT, and Solution 3 (0.2% Uric acid in DI water) for 20 minutes at RT. Between Solution 2 and 3, extensive washes with sterile water (2 washes, 5 minutes each) were given. Samples were dehydrated using an ethanol gradient (25%, 50%, 75%, 95%, and 100%) with two 10-minute incubations at each concentration. Prepared samples were loaded onto SEM stubs and stored in a vacuum until imaging. Samples were sputtered with gold (Jeol Smart Coater) and imaged on the Ultra55 FE-SEM Carl Zeiss mono scanning electron microscope at the Micro Nano Characterization Facility (MNCF), Centre for Nanoscience, Indian Institute of Science.

### Fluorescence microscopy of fixed samples

Images of fixed samples on coverslips were acquired using Total Internal Reflection Fluorescence (TIRF) microscopy using the Nikon Ti2 Eclipse equipped with solid-state lasers (488 nm, 561 nm, and 647 nm were used), 100X TIRF objective (NA=1.49, Apo TIRF 100X Oil DIC N2, Nikon), and a DS-Qi2 CMOS camera (Nikon). Structured Illumination super-resolution Microscopy (SIM) images of fixed Jurkat synapses were acquired on a Zeiss Elyra PS.1 system equipped with a 100X objective (NA=1.46, alpha Plan-Apochromat Oil DIC M27, Zeiss), widefield illumination with laser (488 nm was used), and Z-stacks were acquired for each cell with a slice interval of 0.2 µm.

### Live imaging with ROI-based photoirradiation

Live imaging of cells with laser-based photoirradiation and FRAP was conducted on an Olympus FV3000 laser scanning confocal microscope with lasers (405 nm, 488 nm, 561 nm, 640nm), using the 20X objective (NA=0.75, Air, UPLSAPO 20, Olympus) for T-cell migration assays and the 100X objective (NA=1.4, Oil, UPLSAPO 100, Olympus) for subcellular photoirradiation in Jurkats and adherent cell lines. Photoirradiation was also carried out in a Nikon Ti2 TIRF-FRAP system for *C. elegans* (Brightfield illumination and 405 nm laser for photoirradiation) using a 4X objective, and for Jurkat T-cells expressing Arp3-eGFP (TIRF mode and 488 nm laser for bleaching) with a Galvo XY scanning module, and a Hamamatsu camera. A Tokai Hit heating stage (37 °C, 5% CO_2_, humidified) was used during all live cell imaging experiments, and the temperature was maintained at 22 °C for *C. elegans* experiments. For on-stage photocrosslinking of AZ at sub-cellular or population-level photoirradiation in the Olympus system, a 405 nm laser was used within a defined ROI with average irradiance per ROI of 800 W/cm^2^ for 5 seconds in a sub-cellular setup, and average irradiance per ROI of 4.1 W/cm^2^ for 10 seconds in a multi-cell setup. At the organism-level photoirradiation in the Nikon system, a 405 nm laser was used within a defined ROI with an average irradiance per ROI of 8 W/cm^2^ for 2 seconds.

#### Subcellular photoinhibition of actin nucleation in Jurkat T-cells

Glass-bottomed 8-well chambers (Labtek) were coated overnight at 37 °C with 1 µg/mL His-ICAM-1 (Sinobiological) and 5 µg/mL of anti-CD3 (clone OKT3—human) (BioXCell). Wells were washed with warm PBS before Jurkat T-cells stably expressing Lifeact-Citrine, or transduced with F-Tractin-mCherry lentivirus, at a density of 0.1 million cells in 150-200 µL imaging media per well, were allowed to synapse for 5 minutes, after which AZ at a concentration of 5 µM, CK666 at 50 µM, or control media was added. Circular ROIs were drawn in branched actin-rich peripheral regions of the cell. Imaging was carried out for up to 3 minutes per cell in continuous mode (frame interval of 0.88 seconds), with a photoirradiation of 5 seconds with the 405 nm laser after the first 30 seconds.

#### Subcellular photoinhibition of actin nucleation in adherent cell lines

HEK293T and CHO cells transduced with F-Tractin-mCherry lentivirus were seeded at 15,000 cells per well in an 8-well chamber (Labtek), at 37 °C overnight. AZ was added at a concentration of 5 µM, and CK666 at 50 µM,, and imaging was started immediately upon addition for 75 seconds in total with a 2-second frame interval, and a 5-second photoirradiation with the 405 nm laser after the first 10 seconds was performed in circular ROIs in peripheral regions.

#### Fluorescence recovery after photobleaching (FRAP) of Arp3-eGFP

HEK293T and CHO cells were seeded at 20,000 cells per well in an 8-well chamber (Labtek) at 37 °C overnight. Cells were transfected in the chamber with Arp3-eGFP the day before imaging. Imaging for each cell was carried out for 75 seconds in total with a 2-second frame interval and a 2-second photobleaching with the 488 nm laser within a circular ROI after the first 10 seconds. For Jurkats nucleofected with Arp3-eGFP, cells were seeded at approximately 0.2-0.3 million viable cells in 100 µL media per well of an 18-well chamber (Ibidi) coated for 3-hours with 1 µg/mL His-ICAM-1 (Sinobiological) and 5 µg/mL of anti-CD3 (clone OKT3—human) (BioXCell). Each well was imaged for up to 1 hour, with imaging for each cell carried out for 70 seconds in total with a 0.8-second frame interval and a 0.5-second photobleaching with the 488 nm laser within a circular ROI placed peripherally after the first 10 seconds.

#### Migration assays with population level photoirradiation

18-well chambers (Ibidi) were coated at 37 °C overnight with 2 µg/mL His-ICAM-1 (Sinobiological) and 2 µg/mL anti-CD3 functional antibody (clone 2C11—mouse) (BioXCell). Wells were washed with warm PBS before primary T-cells were added at 0.8 million cells per well in 200 µL imaging media. Cells were allowed to synapse for 5 minutes, then washed twice with warm PBS to remove unbound cells, and imaging media containing 5 µM AZ was added. Brightfield imaging was carried out for 10-15 minutes in total with a 10-second frame interval and a 10-second photoirradiation with the 405 nm laser at the start, within a rectangular ROI encompassing half the field of cells.

### *C. elegans* (organism level) photoirradiation

The *Caenorhabditis elegans* Bristol N2 strain was used as the wild-type reference in all experiments. Worms were maintained following standard procedures on nematode growth medium (NGM) plates seeded with Escherichia coli OP50 as the food source ^58^. Cultures were grown at 22 °C unless otherwise specified. Age-synchronized populations were obtained using standard alkaline hypochlorite treatment. Briefly, gravid adults were treated with a bleaching solution containing commercial household bleach and 5 N NaOH to isolate eggs ^59^. The recovered eggs were washed and transferred onto fresh NGM plates to allow synchronous development to the desired stage. For the preparation of imaging plates, an overnight culture of E. *coli* OP50 was heat-killed by incubating it at 90 °C for 1 hour. After heating, the culture was allowed to cool to room temperature. The resulting heat-killed suspension was then used to seed behavioural assay plates, preventing any effects of active bacterial metabolism. 50 µL of AZ diluted in sterile water to 1 mM (16.9 µg) for AZ for whole-field illumination and on-stage photoirradiation, respectively, was spread with a sterile spreader and dried completely on 30 mm dishes with 2 mL of solidified, freshly prepared NGM. Control dishes contained equivalent amounts of DMSO. 20 µL of heat-killed OP50 was added to the centre of the dish and dried completely. Age-synchronized L4-stage worms were transferred to the plates and incubated in the dark for 20 minutes.

For whole-field illumination, worm locomotion recordings were acquired using an IR wavelength in a WormLab imaging setup for 45 minutes under 488 nm LED illumination, and controls were similarly imaged in the absence of LED. Worm velocity was quantified using MBF WormLab software at t=0, 10, 15, and 20 minutes. For on-stage photoirradiation, a 405 nm laser was used to photostimulate for 5 seconds at the head region of the worm, up to 3 pulses per worm, with quantification performed after each round of pulses. Worm movement was monitored immediately after photoirradiation under 4X magnification, brightfield illumination, at a 0.4-second frame interval for up to 1 minute. After a 1-hour rest period at 22 °C, worms were photostimulated once per worm under the same conditions and imaged again to assess long-term effects.

### Computational simulation

Brownian dynamics computational simulations of the actin cytoskeleton were performed, as described in prior studies ^41,60,61^. Briefly, actin filaments can undergo stochastic nucleation, polymerization, and depolymerization with rates of k_N [μM^-1^ s^-1^], k_P [μM^-1^ s^-1^], and k_D [s^-1^], respectively. In these simulations, to reduce complexity, a fixed ratio of k_N: k_P: k_D = 1: 300: 300 was maintained at baseline in order to maintain comparable average actin filament lengths (∼400 to 450 nm) and average number of actin filaments (∼2900) between conditions. The simulation domain was 8 x 8 x 0.5 μm^3^ to mimic flat cell regions or the actin cortex. The actin concentration was 25 μM. Geometric and concentration parameters are chosen to be at comparable scales to living cells while keeping the computational cost reasonable.

### Image analysis

Image analysis was carried out in Fiji ImageJ software. For total internal reflection fluorescence (TIRF) microscopy images, regions of interest (ROIs) were created for cells by thresholding and detection of particles, and mean intensity was measured for all channels. For structured illumination microscopy (SIM) images of fixed Jurkat synapses, three slices from the z-stack encompassing the synapse plane were averaged for each cell. High-intensity features of the cytoskeleton in TIRF and SIM microscopy images were extracted and estimated as described previously ^41,62^. Briefly, a Gaussian filtered image was created and then subtracted from the raw image to create the “extracted features” image, which highlights denser points in the actin cytoskeleton, such as junctions in the peripheral actin network and actin foci, as described previously ^62^. To segment the synapse, thresholding-based detection was used to create an ROI for the whole cell and a donut-shaped ROI of just the periphery (lamella + lamellipodia). The whole cell ROIs were scaled down to 80% of their size (outer 20% was designated as the lamellipodial region ^38^) to encompass the lamella and the central region of the synapses, and integrated density was measured within the individual regions.

For scanning electron microscopy (SEM) images of the cytoskeleton in Jurkat T-cell synapses, contrast of features was enhanced equally across conditions, and all images were subdivided into six regions. Regions containing empty coverslip outside of the cell or nuclear regions were removed. Images were loaded into Ilastik (version 1.4.1), and pixels were characterized into three features: dense features, fine features, and gaps. Probability maps of these features were exported and opened in Fiji, thresholded to convert to binary masks, and the fraction of each feature was computed from the histogram. For network analysis, the ‘Ridge Detection’ plugin in Fiji was used to detect the cytoskeleton as ridges, which were then converted to a mask and skeletonized using the ‘Skeletonize 2D/3D’ plugin. Skeletons were analysed to extract the number of branches and junctions per image. “Trees” were defined as 4 or more linked branches. Trees, branches, and junctions per region were divided by region area to compute density before plotting.

Migrating T-cells and *C. elegans* were tracked manually on Fiji, and values of X and Y coordinates were used to plot individual cell trajectories using custom codes in MATLAB (R2023b) (The MathWorks Inc.) software. For comparison between photo-inhibited and non-inhibited halves of the field of view in migrating T-cells, distance values were used to compute mean speed and total distance travelled, and the time spent stationary by cells was computed from the fraction of times the instantaneous velocity values became zero. Similarly, for worms, distance values were used to compute mean speed and total distance travelled.

Fluorescence recovery after photobleaching (FRAP) analysis of F-Tractin-mCherry and Arp3-eGFP signal in HEK293T, CHO, and Jurkats was performed in Fiji using the ‘multi-measure’ function to record mean intensity within an ROI of a cell over time. Values from each ROI were normalized to the pre-FRAP values, and average curves were plotted ± SEM over time. Recovery curves were fitted using a single-exponential model in OriginPro 2024 (OriginLab). Simulation videos were analysed in Fiji, where a 150X300 pixel rectangular ROI was used to measure mean intensity in the actin-depleted/bleached and distal zones over time using the multi-measure function. The bleached region was divided by distal region values to retrieve the normalized intensity value over time for each condition.

### Statistical analysis

All statistical analyses were performed using GraphPad Prism 5 software (Version 5.03) for Windows. For comparison of 2 groups, non-parametric (Mann-Whitney) two-tailed t-tests were performed. For comparison of more than 2 groups, the Kruskal-Wallis test with Dunn’s multiple comparisons test was performed. For the proliferation assay, treated groups were compared to the untreated control using a one-way ANOVA with Dunnett’s post-hoc multiple comparison test. For the pulse-wise worm migration speed, whole-field behavioural assay, two conditions were compared with a 2-way mixed-model repeated measures ANOVA. For F-Tractin-mCherry recovery curves, curves were compared using a 2-way repeated measures ANOVA with Dunnett’s post-hoc multiple comparison test.

## Acknowledgements

We thank the Biological Sciences Divisional Microscopy Facility, Divisional Flow Cytometry Facility, and Central Animal Facility at the Indian Institute of Science. We thank Eric Dane for valuable discussions. SM acknowledges GM, AM, and IJ for analysis-related discussions and the Prime Minister’s Research Fellowship (Graduate fellowship). We thank the Council of Scientific and Industrial Research fellowships for AC and PAS; SAA acknowledges support from ANRF (ANRF/ARG/2025/005174/CS); SAA acknowledges support from ICMR (IIRPSG-2025-01-06141). SK acknowledges generous support from an Intermediate Fellowship from the India Alliance DBT-Wellcome Trust (IA/I/23/1/506757). A patent on Azido-CK666 synthesis (Patent #579486, Application #202441043403) has been granted to SM, SAA, ED, and SK.

## Notes

### Competing Interest Statement

The authors have declared no competing interest.

