## Supplemental Text for "An optopharmacological tool for on-demand perturbation of branched actin polymerization"

**Supplemental data**

**Supplementary Figure 1: Purification and chemical analysis of AZ**

(a-b) High-resolution mass spectrometry (HRMS) analysis during synthesis and purification of AZ. Plots show counts plotted against mass/charge peaks showing (a) the intermediate product and (b) the final product (AZ). (c-d) $^{1}$H-NMR spectra recorded for the intermediate and final products of AZ synthesis, with characteristic peaks confirming their chemical structures. (c) Intermediate product ($^{1}$H NMR (CDCl3, 400 MHz): $\delta$(ppm) 6.5-8.5 (7H, Ar-H), 7.8 (N-H, s), 6.69 (N-H, s), 3.67 (2H, t), 2.9 (2H, t), 2.34 (3H, s).) (d) and AZ ($^{1}$H NMR (CDCl3, 400 MHz): $\delta$(ppm) 6.5-8.5 (7H, Ar-H), 7.9 (N-H, s), 6.9 (N-H,s) 3.67 (2H, t), 2.9 (2H, t), 2.34 (3H, s)). (e) *In-vitro* crosslinking of dimethylamine with AZ in the absence of light, with reactants and products analyzed using LCMS—UV chromatogram (absorbance plotted against time) and intensity plotted against mass/charge peaks (inset graph).

**Supplementary Figure 2: Irradiation time optimization and cell health**

(a) Histogram of CFSE dilution peaks of murine primary T-cells obtained using flow cytometry. Peaks are indicative of generations/divisions. (b) Graph showing the percentage of cells divided within 48 hours under the indicated irradiation (254 nm) conditions, compared to the unirradiated control. Each point in the graph represents mean values derived from a single replicate ± Standard Error of Mean (n= 3 samples analyzed per condition). $^{ns}P \geq 0.05$, $^{*}P < 0.05$, $^{***}P \leq 0.001$, as determined using a Kruskal-Wallis test with Dunn's multiple comparisons test. N = 2 replicates with similar results.

**Supplementary Figure 3: Characterization of AZ activity *in-cellulo***

(a) TIRF microscopy images showing levels of F-actin (phalloidin) and phospho-HS1 (pHS1) at the murine primary T-cell synaptic interface. "CON", non-drug control cells; "AZ", AZ-treated cells; "+ or - Light", presence or absence of photoirradiation. Image scale bar = 10 µm. (b) Graphs comparing the mean intensity of F-actin and pHS1 in raw and processed (extracted features) images. Each point in the graph represents mean values derived from a single cell, further normalized to "CON – Light" ± Standard Error of Mean (n= 50-60 cells analyzed per condition). $^{ns}P \geq 0.05$, $^{***}P \leq 0.001$, $^{****}P \leq 0.0001$, as determined using a Kruskal-Wallis test with Dunn's multiple comparisons test. N = 2 replicates with similar results. (c) TIRF microscopy images showing levels of Arp2/3 complex (by Arp2 antibody staining) at the Jurkat T-cell synaptic interface. “CON”, non-drug control cells; “AZ”, AZ-treated cells; “+ or - Light”, presence or absence of photoirradiation. Image scale bar = 10 μm. (d) Graphs comparing the mean intensity of Arp2 in raw images (top) and processed (extracted features) (bottom). Each point in the graph represents mean values derived from a single cell, further normalized to “CON + Light” ± Standard Error of Mean (n= 22-39 cells analyzed per condition). $^{ns}P \geq 0.05$, $^{**}P \leq 0.01$, $^{***}P \leq 0.001$, $^{****}P \leq 0.0001$, as determined using a Kruskal-Wallis test with Dunn’s multiple comparisons test. N = 2 replicates with similar results.

**Supplementary Figure 4: Characterization of AZ reversibility**

(a) TIRF microscopy images showing levels of F-actin (phalloidin) at the Jurkat T-cell immunological synapse. “CON”, non-drug control cells; “AZ”, AZ-treated cells; “+ or - Light”, presence or absence of photoirradiation; “WO”, ‘Wash Out’ condition. (b) Graphs comparing the mean intensity of F-actin (phalloidin) in raw and processed (extracted features) images. Each point in the graph represents mean values derived from a single cell, further normalized to “CON – Light” ± Standard Error of Mean (n= 50-69 cells analyzed per condition). $^{****}P \leq 0.0001$, as determined using a Kruskal-Wallis test with Dunn’s multiple comparisons test. N = 3 replicates with similar results. (c) TIRF microscopy images showing levels of F-actin (phalloidin) at the primary T-cell immunological synapse from a CK666 washout experiment. “CON”, non-drug control cells; “CK”, CK666-treated cells; “+ or - Light”, presence or absence of photoirradiation; “WO”, ‘Wash Out’ condition. Scale bar = 10 μm. (d) Graphs comparing the mean intensity of F-actin (phalloidin) in raw and processed (extracted features) images. Each point in the graph represents mean values derived from a single cell, further normalized to “CON – Light” ± Standard Error of Mean (n= 80-97 cells analyzed per condition). $^{ns}P \geq 0.05$, $^{****}P \leq 0.0001$, as determined using a Kruskal-Wallis test with Dunn’s multiple comparisons test. N = 3 replicates with similar results.

**Supplementary Figure 5: Characterization of the effects of AZ treatment on the cell cortex using high-resolution imaging**

(a) A schematic showing a typical Jurkat T-cell immunological synapse segmented into radial zones for spatial analysis of cortical actin cytoskeletal structures. (b) Structured Illumination Microscopy (SIM) images of F-actin (phalloidin) in the Jurkat T-cell synaptic plane, comparing the radial subsynaptic distribution of F-actin in raw and processed (extracted features) images. “CON”, non-drug control cells; “AZ”, AZ-treated cells; “+ or - Light”, presence or absence of photoirradiation. Image scale bar = 10 µm; Inset scale bar = 2 µm. (c) Graphs comparing the intensity of extracted F-actin features in the lamellipodia, lamella, and total synaptic interface. Each point in the graph represents mean values derived from a single cell, further normalized to “CON – Light” ± Standard Error of Mean (n= 11-14 cells analyzed per condition). $^{ns}P \geq 0.05$, $^{*}P < 0.05$, $^{**}P \leq 0.01$, as determined using a Kruskal-Wallis test with Dunn’s multiple comparisons test. N = 3 replicates with similar results. (d) Graph compares cell areas in the synaptic plane from SIM images. Each bar in the graph represents mean values derived from single cells, further normalized to “CON – Light” ± Standard Error of Mean (n= 11-14 cells analyzed per condition). $^{ns}P \geq 0.05$, as determined using a Kruskal-Wallis test with Dunn’s multiple comparisons test. N = 3 replicates with similar results. (e) Scanning Electron Microscopy (SEM) images of detergent-extracted Jurkat T-cell immunological synapses. “CON”, non-drug control cells; “AZ”, AZ-treated cells. Inset 1 is a focused SEM image of the cytoskeleton from the region specified in the Whole Cell Extract. A specified portion in Inset 1 is magnified to show finer details in Inset 2 (cytoskeleton) and skeleton masks of detected ridges in Inset 2 (detected ridges). Image scale bar = 2 µm; Inset 1 and 2 scale bar = 200 nm. (f) Graphs comparing density of “trees” (total “trees”/area; left graph), branch density (number of branches/area; middle graph), and junction density (number of junctions/area; right graph). Each point in the graph represents mean values derived from a single analyzed area of cytoskeleton ± Standard Error of Mean (n= 8-9 areas analyzed per condition). $^{**}P \leq 0.01$, as determined using a Mann-Whitney U Test. N = 2 replicates with similar results.

**Supplementary Figure 6: Scalable spatiotemporal perturbation with AZ**

(a) Snapshots from live recordings of Jurkat T-cells in immunological synapse expressing the fluorescent actin probe Lifeact-citrine, showing acute morphological and symmetry changes after localized actin photoinhibition. Jurkat T-cells are treated with 5 µM AZ (present in media) coupled with on-stage photoirradiation within an ROI in the peripheral actin ring with a 405 nm laser. The yellow dotted circle indicates the ROI of photoirradiation (also marked with a lightning bolt). Scale bar = 10 µm. N = 3 replicates with similar results. (b) Trajectories of migrating primary T-cells under control/no-drug conditions with half the field photoirradiated with 405 nm laser. “CON-Light”, non-irradiated cells; “CON+Light”, irradiated cells. (c) Graphs show parameters quantified from the migration trajectories—mean speed, total distance traveled, and percentage of total time spent in the stationary phase. Each point in the graph represents mean values derived from a single cell ± Standard Error of Mean (n= 53-64 cells analyzed per condition). $^{ns}P \geq 0.05$, as determined using a Mann-Whitney U Test. N = 2 replicates with similar results. (d) Acute and chronic (1-hour post-irradiation) effects of on-stage 405 nm laser pulse-wise on *C. elegans*. “Control”, non-drug/control organisms; “AZ”, AZ-treated organisms. Plots compare the mean speed of the head in μm/sec post-photoirradiation, after 1, 2, and 3 pulses per worm within the span of 10 minutes, and after 1 hour rest time. Each point in the graph represents mean values derived from a measurement from a single worm after 1, 2, or 3 pulses of 405 nm laser photoirradiation ± Standard Error of Mean (n= 8 worms per condition). ****$P \leq 0.0001$, as determined using Mann-Whitney U-Tests. N = 2 replicates with similar results. (e) The plot compares the effects of AZ with and without light on *C. elegans* under whole-field (whole organism) illumination conditions. Each point in each curve in the graph represents mean values derived from single worms ± Standard Error of Mean (n= 8-10 worms per condition). Curves are normalized to the respective starting value. **$P \leq 0.01$ at t = 20 min, as determined using a 2-way mixed-model repeated measures ANOVA. N = 2 replicates with similar results.

**Supplementary Figure 7: Actin and Arp2/3 complex fluorescence recovery in different cell lines**

(a) Snapshots from live recordings of localized photoirradiation under control/no-drug conditions using a 405 nm laser in Jurkat T-cells forming an immunological synapse (upper panel), CHO cells (middle panel), and HEK293T cells (lower panel) expressing F-Tractin-mCherry. Yellow circle, ROI used for photoirradiation and measurement of fluorescence recovery post irradiation; white squares, regions magnified in the insets. Image scale bar = 10 μm; Inset scale bar = 2 μm. Kymographs on the right, generated from the slanted dotted lines in the leftmost snapshots, show the local variation in F-Tractin signal over time. (b) Graphs comparing F-actin (F-Tractin-mCherry) signal between Control (“CON”) and CK666-treated (“CK”) conditions in Jurkats, CHO, and HEK293T cells. Each bar represents mean values derived from single cells ± Standard Error of Mean (n= 12-21 cells analyzed per condition). $^{ns}P \geq 0.05$, *$P < 0.05$, as determined using a Mann-Whitney U Test. N = 3 replicates with similar results. (c) Snapshots from live recordings of fluorescence recovery after photobleaching of Arp3-eGFP in Jurkats (upper panel), CHO (middle panel), and HEK293T (lower panel) cells. The yellow circle indicates the ROI used for photobleaching and measurement of fluorescence recovery. Image scale bar = 10 μm. (d) Graphs showing normalized fluorescence recovery of Arp3-eGFP signal within the ROI of Jurkats, CHO, and HEK293T cells, plotted over time. N = 3 replicates with similar results per cell line. Each curve in the graphs represents mean values derived from a single ROI over time, normalized to pre-bleach values ± Standard Error of Mean (1 ROI = 1 curve. n= 30-40 cells per condition) (e) Fitting of fluorescence recovery curves to the single-exponential equation: $Y= Y0 + A*e^{-kX}$; Y= Fluorescence intensity, X= Time, Y0= initial value, k= rate constant, A= Plateau value; $t_{1/2}$= half-life of recovery calculated as $t_{1/2}= 0.693/k$; $r^2>0.90$ for all fits, where the upper set (Green, Control; Red, AZ-treated) shows fitted F-actin (F-Tractin-mCherry) fluorescence recovery and the lower set shows fitted Arp3-eGFP fluorescence recovery.

### Supplementary Movie Legends

**Supplementary Movie 1:** Acute on-stage subcellular inhibition of actin nucleation using localized photoirradiation and 5 μM AZ (present in media) in a LifeAct-Citrine expressing Jurkat T-cell forming an immunological synapse. Oval ROI indicates the region illuminated with a 5-second 405 nm laser pulse. Scale bar = 10 μm; Movie frame rate = 20 FPS.

**Supplementary Movie 2:** On-stage localized subcellular photoirradiation (control or "no drug") in a LifeAct-Citrine expressing Jurkat T-cell forming an immunological synapse. Oval ROI indicates the region illuminated with a 5-second 405 nm laser pulse. Scale bar = 10 μm; Movie frame rate = 20 FPS.

**Supplementary Movie 3:** Three examples of cells exhibiting symmetry change upon acute and subcellular inhibition of actin nucleation using localized photoirradiation and 5 μM AZ (present in media) in LifeAct-Citrine expressing Jurkat T-cells forming immunological synapses. Oval ROIs indicate the regions illuminated with a 5-second 405 nm laser pulse. Scale bar = 10 μm; Movie frame rate = 20 FPS.

**Supplementary Movie 4:** Acute, on-stage inhibition of actin nucleation using localized photoirradiation and 5 μM AZ (present in media) in actively migrating murine primary T-cells. Rectangular ROI indicates the section of the field of view illuminated by a 10-second 405 nm laser pulse immediately before the start of the movie. Cell trajectories are color-coded based on time by a red gradient in the upper section ("AZ + Light") and an orange gradient in the lower section ("AZ – Light"). Scale bar = 50 μm; Movie frame rate = 10 FPS.

**Supplementary Movie 5:** Acute organism-level inhibition of actin nucleation using localized photoirradiation and 1 mM AZ (present in agar plate) or control (no drug/DMSO) in *C. elegans*. The bright moving spot of light appearing in the movie indicates the 2-second 405 nm laser pulse for photoirradiation during imaging. Scale bar = 500 μm; Movie frame rate = 25 FPS.

**Supplementary Movie 6:** Acute organism-level inhibition of actin nucleation using localized photoirradiation and 1 mM AZ (present in agar plate) or control (no drug/DMSO) in *C. elegans* conjugates/pairs. The bright moving spot of light appearing in the movie indicates the 2-second 405 nm laser pulse for photoirradiation during imaging. Scale bar = 500 μm; Movie frame rate = 25 FPS.

**Supplementary Movie 7:** Acute subcellular inhibition of actin nucleation using localized photoirradiation and 5 μM AZ (present in media) in an F-Tractin-mCherry expressing Jurkat T-cell forming an immunological synapse (left), CHO cell (middle), and HEK293T cell (right). Circular ROI indicates the irradiated region (5 seconds, 405 nm laser). Scale bar = 10 μm; Movie frame rate for Jurkats = 10 FPS; Movie frame rate for CHO and HEK293T= 5 FPS.

**Supplementary Movie 8:** On-stage localized subcellular photoirradiation (control or “no drug”) in an F-Tractin-mCherry expressing Jurkat T-cell forming an immunological synapse (left), CHO cell (middle), and HEK293T cell (right). Circular ROI indicates the irradiated region (5 seconds, 405 nm laser). Scale bar = 10 μm; Movie frame rate for Jurkats = 10 FPS; Movie frame rate for CHO and HEK293T= 5 FPS.

**Supplementary Movie 9:** Fluorescence Recovery After Photobleaching (FRAP) of Arp3-eGFP expressing Jurkat T-cell in an immunological synapse (left), CHO cell (middle), and HEK293T cell (right). Circular ROI indicates the bleached region. Scale bar = 10 μm; Movie frame rate for Jurkats = 10 FPS; Movie frame rate for CHO and HEK293T= 5 FPS.

**Supplementary Movie 10:** Square simulation space showing local disappearance (localized nucleation inhibition) and recovery of actin filaments. F-actin filaments (cyan), actin-crosslinking proteins or ACPs (orange dots), a region representative of localized photoinhibition (purple rectangle), and a region representative of the surrounding non-inhibited space (dotted rectangle) are shown, with expressions below describing the baseline nucleation rate ($k_{N,\ Baseline}$) in terms of ROI nucleation rate ($k_{N,\ ROI}$).
